# Relative evolutionary rate inference in HyPhy with LEISR

**DOI:** 10.1101/206011

**Authors:** Stephanie J. Spielman, Sergei L. Kosakovsky Pond

## Abstract

We introduce LEISR (*Likehood Estimation of Individual Site Rates*, pronounced “laser”), a tool to infer relative evolutionary rates from protein and nucleotide data, implemented in `HyPhy`. `LEISR` is based on the popular `Rate4Site` (Pupko et al., 2002) approach for inferring relative site-wise evolutionary rates, primarily from protein data. We extend the original method for more general use in several key ways: i) We increase the support for nucleotide data with additional models, ii) We allow for datasets of arbitrary size, iii) We support analysis of site-partitioned datasets to correct for the presence of recombination breakpoints, and iv) We implemented LEISR as MPI-enabled to support rapid, high-throughput analysis. LEISR is available in HyPhy starting with version 2.3.8.

## INTRODUCTION

Evolutionary rate inference is a fundamental analysis in computational molecular evolution (Echave et al., 2016). A widely-used tool for inferring evolutionary rates from phylogenetic protein data is `Rate4Site`, which exists both as a server and a command-line tool (Pupko et al., 2002). Although this method has proven extremely useful over the years, garnering nearly 500 citations, Rate4Site has several limitations: i) it cannot analyze more than ~ 200 – 300 sequences because of numerical underflow issues; ii) it often fails to converge to stable estimates if data are sufficiently complex even with relatively few (25 – 100) sequences; and iii) it accepts primarily protein data only and has limited nucleotide utility. As the number of available genomic sequences continues to rapidly expand, tools to analyze large data sets of any genomic type (protein and nucleotide), are needed.

To this end, we introduce a generalization of `Rate4Site` in the leading molecular evolution inference platform `HyPhy` (version ≥ 2.3.8), which we term “LEISR” (**L**likehood **E**stimation of **I**ndividual **S**ite **Rates**). LEISR can be used to infer relative evolutionary rates from either nucleotide or protein data, thereby providing a flexible and fast platform for rate inference on non-coding data^1^. `LEISR` has been successfully tested with alignments containing up to 10,000 sequences, 2 to 3 orders of magnitude beyond what Rate4Site can fit. In addition, LEISR is additionally `MPI`-enabled to support rapid inference from datasets with many sites, by distributing optimization tasks to multiple compute nodes. Like other methods in `HyPhy`, `LEISR` allows users to provide partitioned alignments, with separate phylogenies for each partition, to correct for the effect of recombination during rate inference. Such partitioned alignments can be obtained, for example, with the method GARD (Kosakovsky Pond et al., 2006) in `HyPhy`.

## APPROACH

`LEISR` proceeds in two steps. It first obtains estimates of alignment-wide branch lengths under a user specified substitution model (Table 1), and it then for each site, it infers a scaler parameter, *r_s_*, that is used to uniformly scale all the branch lengths of the partition-specific tree at the site. *r_s_* can therefore be interpreted as the evolutionary rate at a specific site relative to the alignment-wide mean rate.

**Table 1.** Nucleotide and Protein models, both generalist and specialist, available for use in `LEISR`, as of `HyPhy` version 2.3.8. Future HyPhy versions are expected to include more models. We note that users can define and fit other parametric and empirical models with the use of HBL, the `HyPhy` batch language.

| Data type | Models |
| --- | --- |
| Nucleotide | GTR(Tavare, 1984), HKY85(Hasegawa et al., 1985), JC69(Jukes and Cantor, 1969) |
| Protein, Generalist | LG(Le and Gascuel, 2008), WAG(Whelan and Goldman, 2001), JTT(Jones et al., 1992), JC69(Jukes and Cantor, 1969) |
| Protein, Specialist | mtMam(Yang et al., 1998), mtMet(Le et al., 2017a)(Le et al., 2017b), mtVer(Le et al., 2017b), cpREV (Adachi et al., 2000),gcpREV (Cox and Foster, 2013), HIV B/W (Nickle et al., 2007), AB (Mirsky et al., 2015) |

As this algorithm proceeds, `HyPhy` will write markdown-formatted status-indicators to the console, including the inferred site-wise maximum-likelihood rate estimates with the approximate 95% confidence interval (CI) obtained via profile likelihood. All final output is written to a JSON-formatted file, named as the input data file with the suffix .LEISR.JSON. Site-wise rates are stored in the top-level JSON field MLE, whose content field contains a row for each site’s inferred rates. Individual values in each row correspond to information given in the headers key. We note that a general description of HyPhy output JSON contents is available from http://www.hyphy.org.

Users are free to transform these rates in a manner that suits their given analyses. For example, `Rate4Site` computes a standard score for each site, and other applications have called for normalizing each rate by the average (or median) gene-wide rate (Jack et al., 2016; Sydykova et al., 2017). This latter scheme re-scales the average gene rate as 1, lending a more intuitive interpretation to each site’s rate, i.e. a rate of 2 indicates that a site evolves twice as quickly as does an average site, and a rate of 0.5 indicates that a site evolves half as quickly as does an average site. As empirical rate distributions are generally overdispersed and zero-inflated, we suggest to normalize by the median rather than the mean, should normalization be desired. We note that even raw rate estimates generated by `LEISR` are already defined relative to the jointly-inferred partition mean rate.

`Rate4Site` offers two statistical frameworks, maximum-likelihood (ML) (Pupko et al., 2002) and empirical Bayes (Mayrose et al., 2004), for rate inference, where the latter approach requires a discrete gamma distribution of rates be used during branch length optimization. LEISR, on the other hand, only employs maximum-likelihood, and rate variation (if chosen) during branch length optimization can be modeled either with a discrete gamma distribution (Yang, 1993) or the general discrete distribution (GDD) (Kosakovsky Pond and Muse, 2005). Although we provide the option to employ rate variation for branch length optimization, we encourage users to opt for no rate variation. Indeed, the desired behavior for this method is for *only* the relative site-wise rates to contain information about site-wise evolutionary rate heterogeneity. If branch length optimization considers rate variation, then this information will be “conflated” between these two parameters (branch lengths and site rates). Indeed, one can view `Rate4Site` and `LEISR` as non-parametric rate estimation methods, whereas gamma and GDD are parametric estimation methods, and layering the two would be inefficient.

## RESULTS

We confirmed that LEISR yields comparable inferences to `Rate4Site` using simulations. For each of three random phylogenies with 25, 50, and 100 taxa each, we simulated 10 replicate alignments, each with 100 sites, under the WAG model of protein evolution (Whelan and Goldman, 2001). Our simulations modeled rate heterogeneity among sites with a discrete gamma distribution with 20 categories and a shape parameter of 0.4. We note that each replicate number used the same model parameterizations for all three trees (i.e. replicate 1 employed the same model for 25, 50, and 100 taxa). Simulations were conducted using the simulation library `pyvolve` (Spielman and Wilke, 2015) written in Python.

We then inferred relative evolutionary rates in `LEISR` in two modes: turning off rate heterogeneity during branch length optimization (“LEISR”), and specifying a four-category discrete gamma distribution during during branch length optimization (“LEISR+G”). We again inferred rates in two modes in Rate4Site: without rate heterogeneity during branch length optimization (“R4S”) and with a four-category discrete gamma distribution (“R4S+G”). We conducted all Rate4Site inferences using their maximum-likelihood algorithm (in contrast to their empirical Bayes approach). Note that the default number of rate categories for this step in `Rate4Site` is 16, but `Rate4Site` failed with errors for all 100-taxa simulations. We therefore used four rate categories, to both achieve a fair comparison with LEISR and to ensure that `Rate4Site` could complete inferences. Finally, for each alignment inference, we normalized rate estimates by dividing all rates by the mean site rate estimate, as described earlier.

In Figure 1, we show *r*^2^ values for Pearson’s linear correlation between `LEISR` and `Rate4Site` inferences, computed across all simulations. Figure 2 shows, for a single representative simulation replicate of 100 taxa, the relationship between inferred site rates across different methods and/or parameterization. Overall, these results demonstrate a nearly complete agreement between `LEISR` and `Rate4Site`, with rate inferences showing the closest agreement when the same option for branch length optimization was specified (i.e. turned off or with a discrete gamma distribution). The r^2^ values further increase as the number of taxa increases, although even with 25 taxa the agreement is remarkably high. This is expected because the precision of inference for individual site rates will increase for larger samples (more taxa, Scheffler et al. (2014)). We therefore find that LEISR provides a robust and reliable platform that can be used in the place of `Rate4Site` when dataset size and/or complexity preclude `Rate4Site` use, or when recombination is suspected.

**Figure 1.**
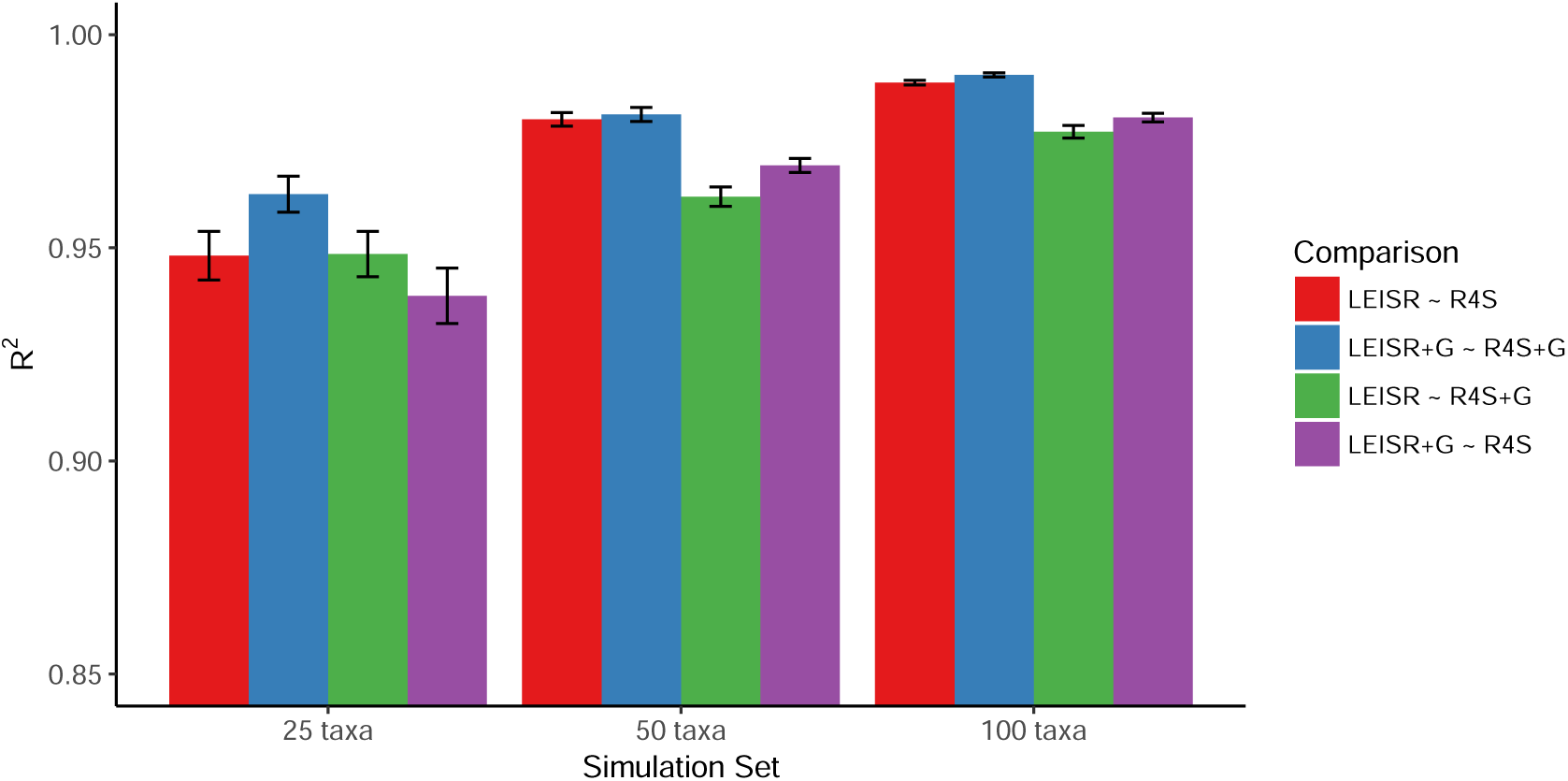
Mean *r*^2^ values (across 10 replicates) between inferred evolutionary rates across platforms and simulations. Bars represent the standard error of the mean. Note that the y-axis of this figure begins at *r*^2^ = 0.85. All code to generate simulations and reproduce figures is available from https://github.com/sjspielman/leisr_validation.

**Figure 2.**
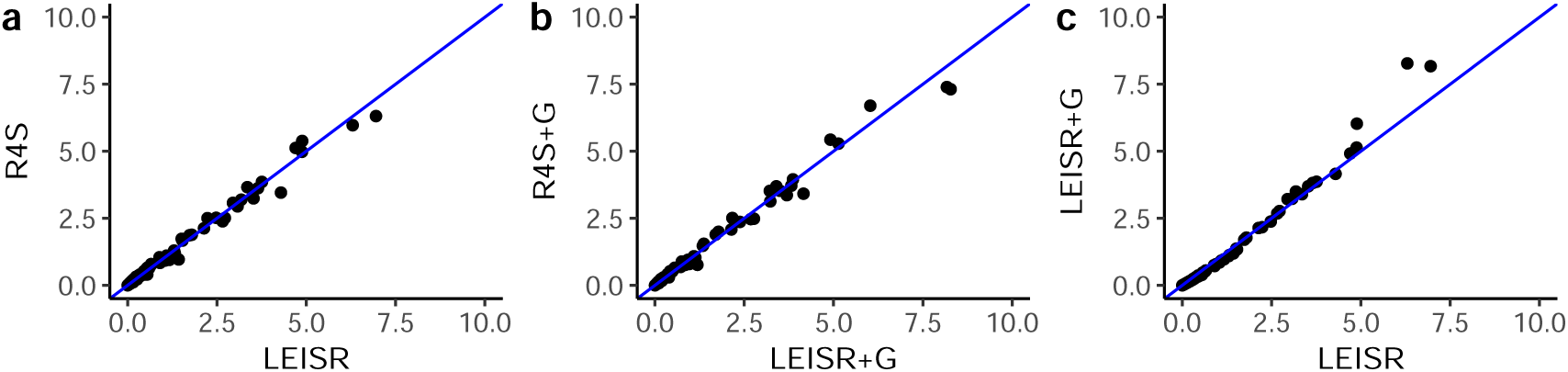
Inferred evolutionary rates for a single simulation replicate with 100 taxa. The line shown in each panel is *y* = *x*. All code to generate simulations and reproduce figures is available from https://github.com/sjspielman/leisr_validation.

## CONCLUSIONS

`LEISR` is accessible as an option in the `HyPhy` analysis menu (“Relative evolutionary rate inference”), which calls the `HyPhy` batchfile `LEISR.bf`, in `HyPhy` version ≥ 2.3.8. We encourage users who employ `LEISR` to additionally cite `Rate4Site` (Pupko et al., 2002), which provides the intellectual basis and historical precedent for our implementation. We note that earlier versions of HyPhy (specifically, ≥ 2.3.6) also contain the LEISR method, although those pre-release versions will have reduced functionality relative to the LEISR implemented in HyPhy version 2.3.8.

## ACKNOWLEDGMENTS

This work was supported in part by grants R01 GM093939 (NIH/NIGMS), R01 AI134384 (NIH/NIAID), and U01 GM110749 (NIH/NIGMS).

